# Sc-TUSV-ext: Single-cell clonal lineage inference from single nucleotide variants (SNV), copy number alterations (CNA) and structural variants (SV)

**DOI:** 10.1101/2023.12.07.570724

**Authors:** Nishat Anjum Bristy, Xuecong Fu, Russell Schwartz

## Abstract

Clonal lineage inference (“tumor phylogenetics”) has become a crucial tool for making sense of somatic evolution processes that underlie cancer development and are increasingly recognized as part of normal tissue growth and aging. The inference of clonal lineage trees from single cell sequence data offers particular promise for revealing processes of somatic evolution in unprecedented detail. However, most such tools are based on fairly restrictive models of the types of mutation events observed in somatic evolution and of the processes by which they develop. The present work seeks to enhance the power and versatility of tools for single-cell lineage reconstruction by making more comprehensive use of the range of molecular variant types by which tumors evolve. We introduce Sc-TUSV-ext, an integer linear programming (ILP) based tumor phylogeny reconstruction method that, for the first time, integrates single nucleotide variants (SNV), copy number alterations (CNA) and structural variations (SV) into clonal lineage reconstruction from single-cell DNA sequencing data. We show on synthetic data that accounting for these variant types collectively leads to improved accuracy in clonal lineage reconstruction relative to prior methods that consider only subsets of the variant types. We further demonstrate the effectiveness on real data in resolving clonal evolution in the presence of multiple variant types, providing a path towards more comprehensive insight into how various forms of somatic mutability collectively shape tissue development.

## 1 Introduction

Somatic evolution, in which cell populations in a tissue accumulate mutations over time, is increasingly recognized as both widespread and important to health and disease (24). This process has traditionally been most intensively studied due to its role in cancer. Cancer develops through an evolutionary process in which an initially healthy somatic cell population accumulates mutations over a series of generations, creating genetic diversity from which fitter, more aggressive cell sub-populations (subclonal populations) expand leading to tumorigenesis and tumor progression (27). Cancers are typically characterized by high rates of mutability and acquisition of diverse forms of mutations, including single nucleotide variants (SNV) , copy number alterations (CNA) and structural variations (SV) (33). More recent studies show that somatic mutability is a constant process in healthy tissues, of which cancer is just one possible endpoint (24), and that similar mutation landscapes play roles in other common diseases, such as neurodegenerative illnesses (25).

Recognition of the importance of somatic evolution in cancers led to the widespread interest in characterizing the evolutionary histories of subclonal populations as a way of understanding mechanisms of disease progression as well as providing guidance towards patient specific treatment plans (4). This has commonly taken the form of computational inference of clonal lineage trees — generally known as tumor phylogenies (3; 31). In these approaches, the evolutionary history of the subclonal populations is modeled as a phylogenetic tree, whose nodes represent the major genetically similar subpopulations in a tumor (subclones) and whose edges represent the accumulation of somatic mutations that occur along the way from the parent to the child. The root of the tree usually represents a putatively healthy common ancestor of all subclones, which has no mutations. Many of the current tumor phylogeny reconstruction methods use bulk DNA sequencing (DNAseq) data collected from short reads of mixtures of cells (11; 9; 7), which are deconvolved to infer the distinct clonal subpopulations and their frequencies (32). While methods for bulk data remain important, the relatively low precision of this deconvolution process has led attention of the field to shift in large part to phylogeny inference from single-cell DNA sequencing (scDNAseq) data (26), which allow one to develop tumor phylogeny methods bypassing the deconvolution step (35). scDNA-seq brings its own challenges, however, including typically high rates of allelic dropouts, false positives, and false negatives due to the lower resolution of the technology (13). While single-cell phylogenetics is in principle a simpler problem than deconvolutional phylogenetics, it is challenging in practice to develop methods robust to scDNA-seq data artifacts.

In part because of this, single cell tumor phylogeny methods have lagged behind bulk methods in the sophistication of the evolutionary models they apply. While the earliest methods for single cell tumor phylogenetics focused primarily or exclusively on CNAs, which could be inferred more robustly from lossy data (26; 34; 6), much subsequent work exclusively relied on SNVs and often with restrictive models of their production. The most commonly used such model for SNVs is the infinite sites model (or perfect phylogeny model) where a mutation can only be gained once and never lost along all the edges of the tree. This model is known to be frequently violated in practice (17) but is also algorithmically convenient (14). SCITE (15) and BSCITE (22) use this model for the phylogeny inference with SNVs. The perfect phylogeny assumption is relaxed in the Dollo or k-Dollo parsimony model, where a mutation can be gained once as before but lost multiple or at most *k* times times respectively, on which SPhyR (10), SASC (1), ConDoR (29) are built. SCARLET (30) improves on the previous methods using a loss-supported model with the combination of Dollo and infinite sites assumptions, incorporating both CNAs and SNVs, and requires the copy number profiles and the copy number phylogeny as input or traverses over all possible trees. In the aforementioned models, a mutation can be gained only once. However, in cancer, parallel evolution is a common event where a mutation can be gained independently in different branches of the tree. SiFit (36) further relax the assumptions of the Dollo model and allows a mutation to be gained multiple times using the finite sites model. A handful of methods exist to combine bulk and single-cell data, including the SNV-centric methods like B-SCITE, PhiSCS (23) and the CNA-centric methods such as Lei et. al. (19) and MEDICC2 (16). While these single-cell tumor phylogeny methods account for CNAs and SNVs, there is no such model for single cells to our knowledge that uses SVs arising from chromosomal instability (CIN), which are often a primary driving force in cancer progression (21).

Here, we introduce the first single-cell tumor phylogeny method, Sc-TUSV-ext, which was developed to provide more accurate and comprehensive single-cell clonal lineage models by performing tree inference using SNVs, CNAs, and SVs simultaneously. The work makes use of ideas previously developed for the bulk sequence methods TUSV (9), which first brought SVs to deconvolutional tree inference; and TUSV-ext (11), which first brought SNVs, CNAs, and SVs together to infer tumor phylogenies from bulk sequencing data. Sc-TUSV-ext uses two-stage approach: 1) perform copy number clustering of single cells using the MEDICC2 distances (16) and hierarchical clustering and 2) extend the ILP frameworks of TUSV (9) and TUSV-ext (11) to infer lineage trees constrained to Dollo parsimony models of SNVs and SVs while approximately minimizing CNA evolution. We show through application to synthetic data and comparison to alternative tools that the more comprehensive Sc-TUSV-ext variant model yields improved accuracy across a variety of mutation parameters and performance measures. We further show it to work effectively on real data and reveal novel insights unavailable to any alternative methods.

## 2 Methods

The goal of our method is to identify single-cell lineage trees from a collection of single-cell genetic variant calls while accounting for SVs, SNVs, and CNAs. Figure 1 gives an overview of the analysis pipeline of Sc-TUSV-ext. At a high level, the method works by first clustering single cells into putative clones, using CNAs as the primary marker to cluster by MEDICC2 distance (16), and subsequently inferring clonal lineage trees (“tumor phylogenies”) for the clones using a model based on the bulk data method TUSV-ext (9; 11) that seeks trees consistent with a Dollo parsimony model of SNVs and SVs so as to minimize an objective balancing minimum copy number distance and consistency of copy number calls across SV breakpoints. Each SNV is represented by a position in the genome and its estimated copy number. We represent each SV as a pair of breakpoints that are found adjacent to each other at the cancer genome but non-adjacent in the reference genome. CNAs are handled by treating the genome as discretized into bins, each of which is assigned a copy number for each single cell or cell cluster.

**Fig. 1.**
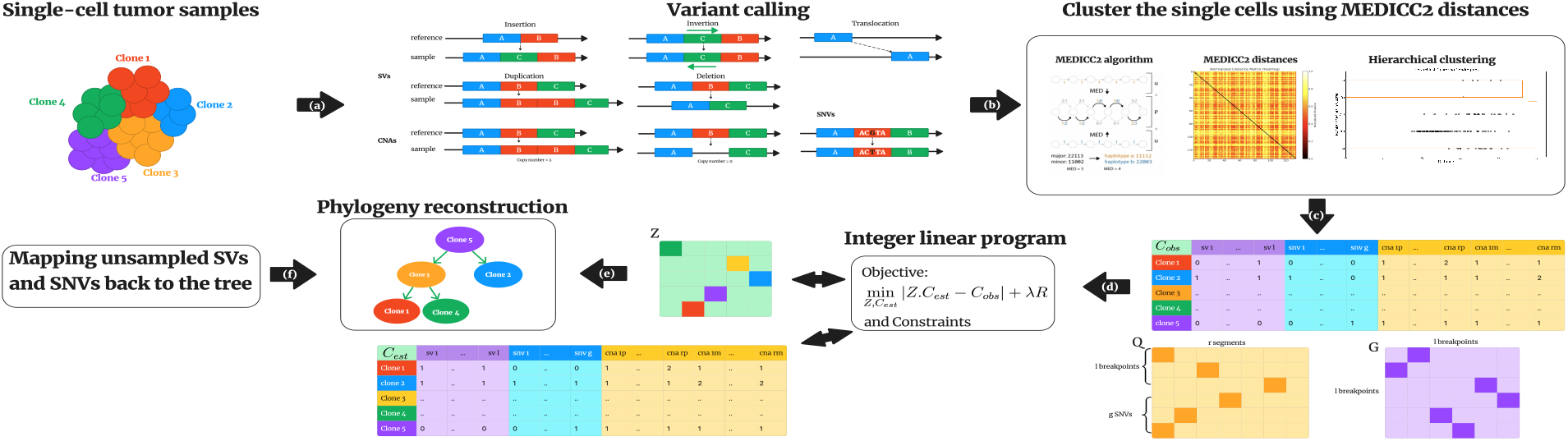
Schematic diagram of the Sc-TUSV-ext method. (a) Starting with the processed variant calls, (b) we cluster the cells using their MEDICC2 distances into a user-defined number of clones and (c) build our variant matrix *𝒞*^obs^, SNVs and breakpoint-to-segment mapping matrix Q, and breakpoint-to-SV mapping matrix G. (d) Then we run our integer linear programming framework and (e) build the phylogeny on a subsampled set of variants. (f) At the end, we map the unsampled SNV and SV breakpoints back to the edges of the phylogeny.

### 2.1 Problem statement

The inputs to Sc-TUSV-ext are processed variant calls of the single cells where we represent SVs with segmental mean copy numbers of paired breakpoint ends, CNAs with allele specific mean copy number for a set of discrete genomic segments, and SNVs with estimated mean copy numbers. Given the processed variant calls for the single cells, we first group *𝒩* single cells into *n* clones based on their MEDICC2 distances. Then, based on the modes copy number information of the single-cells mapped to *n* clones, we build an *n ×* (*l* + *g* + 2*r*) copy number matrix *𝒞*^obs^, *l × r* breakpoint to segment mapping binary matrix *𝒬*, and *l × l* breakpoint-to-SV mapping binary matrix *𝒢*. Given *𝒞*^obs^, *𝒬* and *𝒢*. We fit trees to these data so as to minimize the following objective function

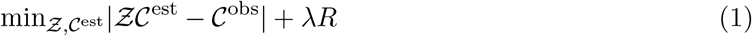

Where *𝒞*^est^ is the *n ×* (*l* + *g* + 2*r*) estimated copy number matrix, 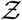 is an *n × n* binary matrix that represents the mapping of clones in the phylogenetic tree according to the assumptions of Sc-TUSV-ext, *R* is the phylogenetic cost, and *λ* is a regularization parameter.

### 2.2 Copy number clustering of single cells using MEDICC2 distance

We first use the MEDICC2 method as described in Kaufmann et. al. (2022) (16) which uses the minimum event distance (MED) for estimating whole genome duplication aware copy number distances from single-cell DNAseq data. We then use hierarchical clustering with complete linkage on the *N × N* MEDICC2 pairwise distance matrix. For the single cells assigned to each cluster, Sc-TUSV-ext uses the modes of the SNV, CNA, and SV calls of cells in the cluster to create the *n ×* (*l* + *g* + 2*r*) *𝒞*^obs^ matrix of cluster representatives.

### 2.3 Coordinate descent algorithm

The bulk of the method is dedicated to solving for a constrained optimization problem, detailed in the subsequent sections. The method is adapted from the earlier TUSV-ext method for building SNV/CNA/SV phylogenies from bulk DNA-seq data (11), itself extended from the earlier CNA/SV TUSV method (9). We describe the full program below but with an emphasis on areas in which Sc-TUSV-ext differs from bulk TUSV-ext.

Our objective function is quadratic and in practice not solvable to optimality within a reasonable time. To resolve this, we follow the earlier TUSV and TUSV-ext in optimizing using a coordinate descent algorithm the poses the problem as two mixed integer linear programs (MILPs) between which we iterate. We solve for 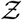 and *𝒞*^est^ iteratively by first solving for 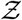 given *𝒞*^est^and *𝒞*^obs^, and then solving for *𝒞*^est^ given 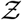 and *𝒞*^obs^. We repeat this process until convergence, when 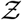 and *𝒞*^est^ does not change for two consecutive iterations, or a predefined maximum number of iterations is reached. We initialize the iteration by shuffling the rows of *𝒞*^obs^and assign them to *𝒞*^est^ then solving the ILP for 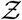.

### 2.4 Estimating 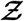 and *𝒞*^est^

Our aim in this step is to assign one clone to each node of the phylogenetic tree. To achieve this, we set the constraints on 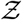 as described in equation 2,3 and 4. We then define the copy number distance portion of the objective in Equation 1 using L1 distance as equation 5, 6 and 7.

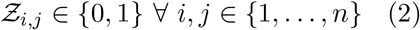

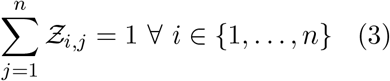

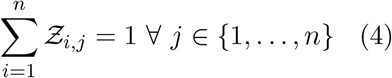

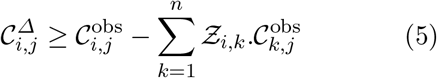

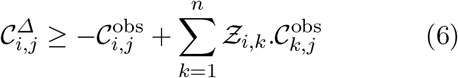

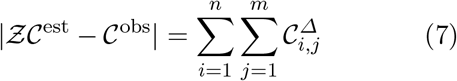

We use these equations to estimate 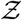 given *𝒞*^obs^ and *𝒞*^est^. Once 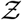 is estimated, we use 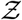, *𝒞*^obs^, *𝒬* and *𝒢* to estimate *𝒞*^est^.

### 2.5 Integer linear programming

#### Phylogenetic and ancestry constraints

As the copy number profiles of the aggregated clones are associated with a underlying phylogenetic tree, we impose phylogenetic constraints on the rows of the *𝒞*^est^matrix as described in Fu et al. (2022) (11), and Eaton et al. (2018) (9). We define an *n × n* edge matrix *E* and *n × n* ancestry matrix *A*, where *n* = 2*n*^*′*^ + 1, to define a binary tree *𝒯* . An edge *e*_*i*,*j*_ = 1 if node *i* is the parent of node *j* and 0 otherwise. Similarly, an entry in the ancestor matrix, *a*_*i*,*j*_ = 1 if node *i* is an ancestor of node *j* and 0 otherwise. For simplicity, we assume that the *n*th row of *𝒞* ^est^ represents the root, rows 1 through *n*^*′*^ represent the leaf nodes and *n*^*′*^ + 1 through *n −* 1 represent the internal nodes. We impose the following constraints on the edge and ancestor matrix:

Root has no incoming edges (Eqn. 8 and leaves have no outgoing edges (Eqn. 9)

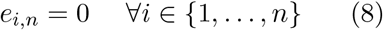

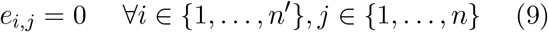

Every node that is not a root has exactly one incoming edge (Eqn. 10) and every internal node has two outgoing edges (Eqn. 11)

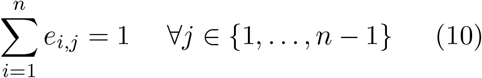

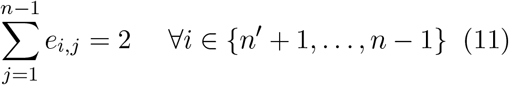

The root is the ancestor of all other nodes (Eqn. 12) and the root has no ancestor (Eqn. 13)

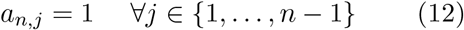

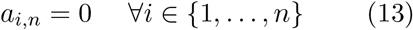

Any node’s parent is its ancestor

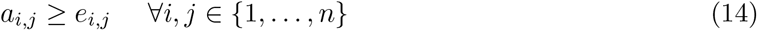

Any child node carries its parent’s ancestors

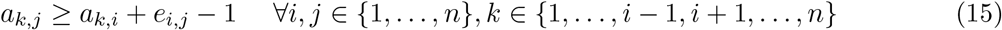

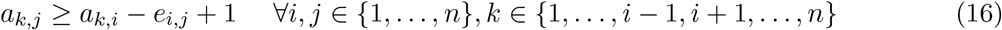

#### Copy number constraints and phylogenetic cost

Similar to TUSV-ext (11), we first impose some constraints on allowed copy numbers. We assume that the root is bi-allelic and has one copy of each segment, no single nucleotide variants, and no breakpoints. We also bound the copy number of each segment to a maximum, *c*_max_. We call the *k, s* th entry of the *𝒞* ^est^ matrix *c*_*k*,*s*_.

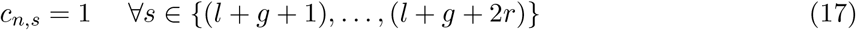

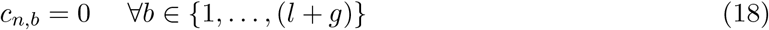

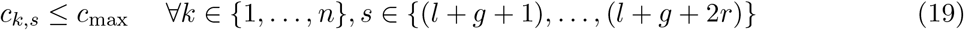

We next model the phylogenetic tree cost as the sum of pairwise L1 distances of the copy numbers across all edges of the tree. While L1 distance is a simplification compared to the MEDICC2 distance used in clustering, it provides an approximate measure of evolutionary distance suitable for prioritizing among otherwise feasible trees. To define this measure as a linear function fo the variables, we follow TUSV-ext in defining two collections of temporary variables 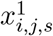 and 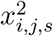, representing the absolute copy number changes across edges *e*_*i*,*j*_.

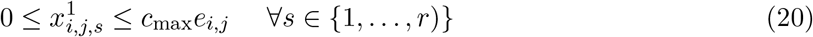

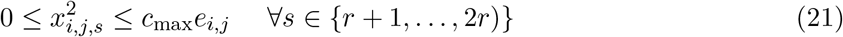

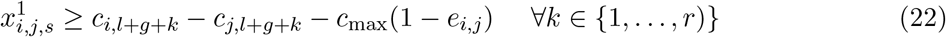

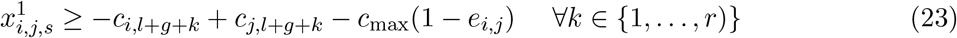

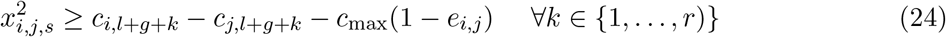

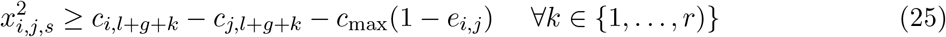

We add the two auxiliary variables for all segments to get the copy number cost, *ρ*_*i*,*j*_ across the edge *e*_*i*,*j*_ (Eqn. 26) and determine the phylogenetic cost R (Eqn. 27).

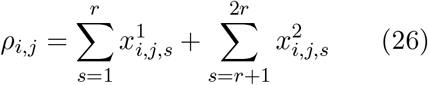

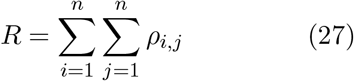

#### Perfect phylogeny on breakpoints and SNVs

Similar to TUSV (Eaton et. al 2018) (9) and TUSV-ext (Fu et. al. 2022) (11), we impose a Dollo phylogeny constraint on SNVs and SVs. For SVs, we require that each breakpoint appears exactly once across one edge of the tree and ensure that a pair of breakpoints appears on a common edge of the tree. We impose these constraints using another binary auxiliary variable, *w*_*i*,*j*,*b*_ corresponding to the *W* ∈ {0, 1}^*n×n×*(*l*+*g*)^ matrix, where *w*_*i*,*j*,*b*_ = 1 if mutation *b* is found across the edge *e*_*i*,*j*_.

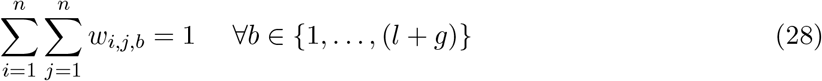

SNVs are treated as a degenerate case of SVs, effectively two breakpoints occupying the same genomic position and also constrained to appear once in the tree. Both kinds of variants can be lost due to deletion concurrent with copy number losses, however, and thus obey Dollo (one appearance but potentially multiple loss) rather than perfect phylogeny (one appearance without loss) constraints.

### 2.6 Mapping of unsampled SNVs and SVs

As the ILP can scale poorly for large number of variants, we apply a heuristic solution of sampling a predefined number of SNVs and pairs of SV breakpoints while keeping all the CNAs, using these to infer the tree topology, and then mapping the unsampled SNVs and SV breakpoints back to the tree. For mapping the unsampled SNVs and SVs, we first create a *n × n* binary matrix *C*^expected^, where each row represents a node and 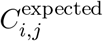 represent the existence of a variant in node j if the variant was introduced in node i , with the perfect phylogeny assumption. For each variant j in our unsampled matrix, *C*^unsampled^, we find the minimum distance node i for the variant and assign variant j to node i (by setting 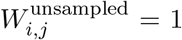). The algorithm is described in finer details in the following pseudocode.

#### Algorithm 1

Map unsampled SNVs and SVs to nodes

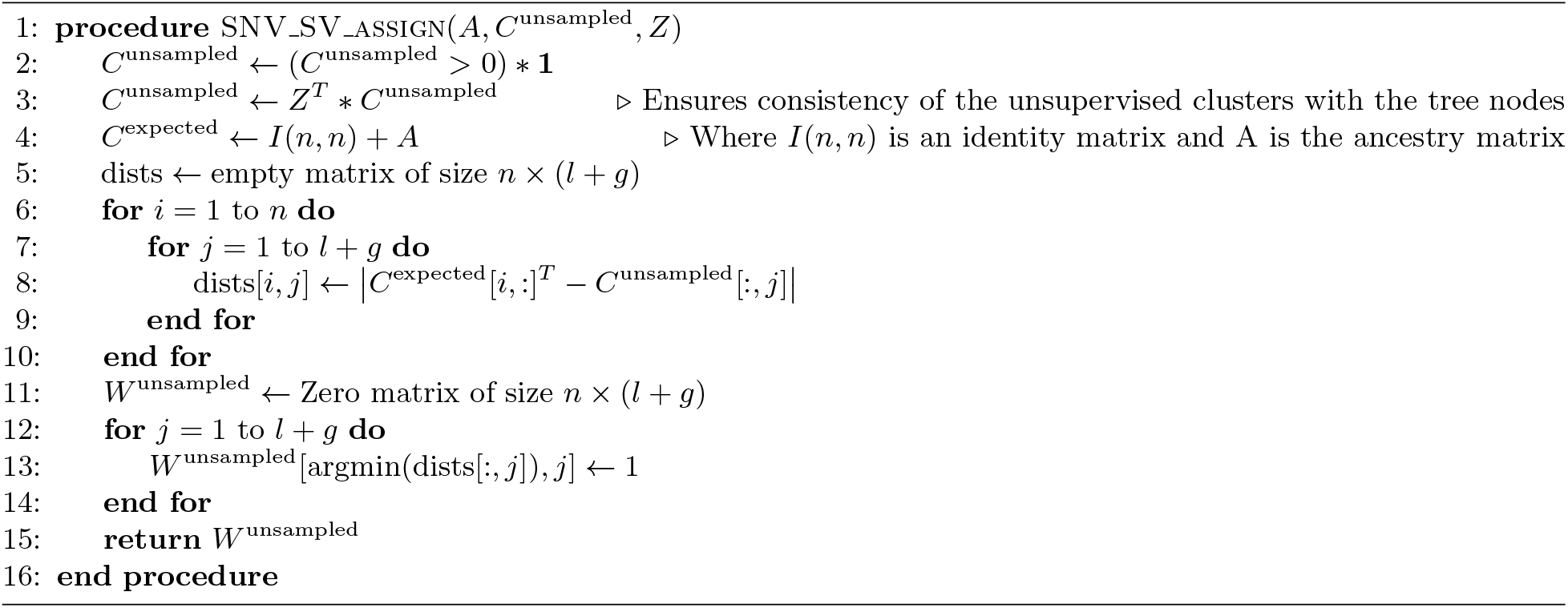

### 2.7 Choice of *λ*

We set the regularization parameter 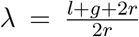, following Fu et al. (2022) (11) and Eaton et al. (2018) (9). We choose the ratio of maximum errors allowed by the first term in our objective function ((*l* + *g* + 2*r*) ∗ *n*) to be equal to the maximum error allowed by the second term (2*r* ∗ *n*).

### 2.8 Implementation

We implemented Sc-TUSV-ext in Python with the licensed gurobi optimizer for solving the integer linear program. All the codes are available at https://github.com/CMUSchwartzLab/Sc-TUSV-ext.

## 3 Results

To establish the effectiveness of Sc-TUSV-ext, we validate it on both simulated and real data and compare it with two other leading methods: SCARLET (30) and MEDICC2 (16). We note that we cannot fully compare Sc-TUSV-ext directly to the other methods, as to our knowledge no other single-cell tumor phylogeny method can build phylogenies incorporating SNV, CNA, and SV data. Most build phylogenies solely on SNV data, potentially accounting for CNAs as a confounding factor in interpreting SNVs. We thus compare to the other methods by comparing Sc-TUSV-ext’s mapping of SNVs alone to that of SCARLET and Sc-TUSV-ext’s phylogenies with MEDICC2’s. While this gives SC-TUSV-ext an advantage over the alternatives, in that it is making use of variant types the other methods cannot use, we believe this provides a fair assessment of the value of considering these variant types collectively, which is the major contribution of Sc-TUSV-ext to this space.

### 3.1 Evaluation on simulated data

We validate the accuracy of our method on simulated data with randomly generated single-cell phylogenies labeled by SNV, SV and CNA mutations. For each simulation, we generate a random binary tree with *n* nodes along with a copy number matix *𝒞*^obs^, where each row represents a single cell and the columns encode SNVs, SVs and CNAs. Then we randomly perturb the copy number profiles to generate the rest of the single cells and construct the clone assignment matrix 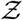 accordingly. We generate each of our simulations for subclones of a single patient on a segment of the whole genome profile that includes chromosome 1 and chromosome 2. We generate the SNVs at uniformly random positions, and SVs sampled from a Poisson distribution with parameter 5,745,000, which is the average length of structural variants observed at the TCGA-BRCA cohort (9). CNAs were generated using a relative probability of 2:1:2:1 for amplifications, inversions, deletions and translocations. To simulate the read counts associated with these genomic alterations, we first modeled the segment-specific read counts using a Poisson distribution with a parameter (mean depth) of 50. We then use these read counts as the trial numbers for a binomial distribution, which generated the variant-associated read counts, with probability derived from the variant allele frequencies.

To test the robustness of our method, we perform a series of experiments including several model conditions — varying number of clones, number of single cells, number of mutations, and error tolerance. For all the simulations, we applied Sc-TUSV-ext with a maximum computational time of 2000 seconds and 5 iterations and 5 random restarts. We ran SCARLET with a setting where the true copy number tree and true copy number profiles for the cells were known — we provided SCARLET with our true simulated phylogeny – and compared the estimated SNVs of SCARLET with SC-TUSV-ext’s. We compared MEDICC2 and Sc-TUSV-ext’s clonal phylogenies with the true phylogenies using the Robinson Fould (RF) distance (28).

### 3.2 Experiment 1: Validation for different level of errors

Given the high noise level typical of single-cell DNA-seq, robustness to data error and loss are important criteria for evaluation. We therefore systematically added x% sequencing error across SNVs and 2*x*% error across allele specific CNAs for each single-cell, where *x* ∈ {0, 2, 4}. For SNV mutations, which are binary, we introduced errors by randomly flipping their values. For CNAs, which are represented as integers in our datasets, we incorporated errors as follows: if a CNA value was 1, it was changed to 0; and conversely, if it was 0 or any integer other than 1, it was changed to 1. For each case, we simulated a 5 clone tree with 100 single-cells, 100 SNVs, 100 SVs and corresponding CNAs. We ran Sc-TUSV-ext with 80 sampled SV breakpoints and 40 SNVs.

Figure 2(g) shows errors in SNV assignments for Sc-TUSV-ext in comparison to SCARLET. Sc-TUSV-ext outperforms SCARLET across error rates, which we attribute to its ability to leverage the additional mutation types unavailable to SCARLET.

**Fig. 2.**
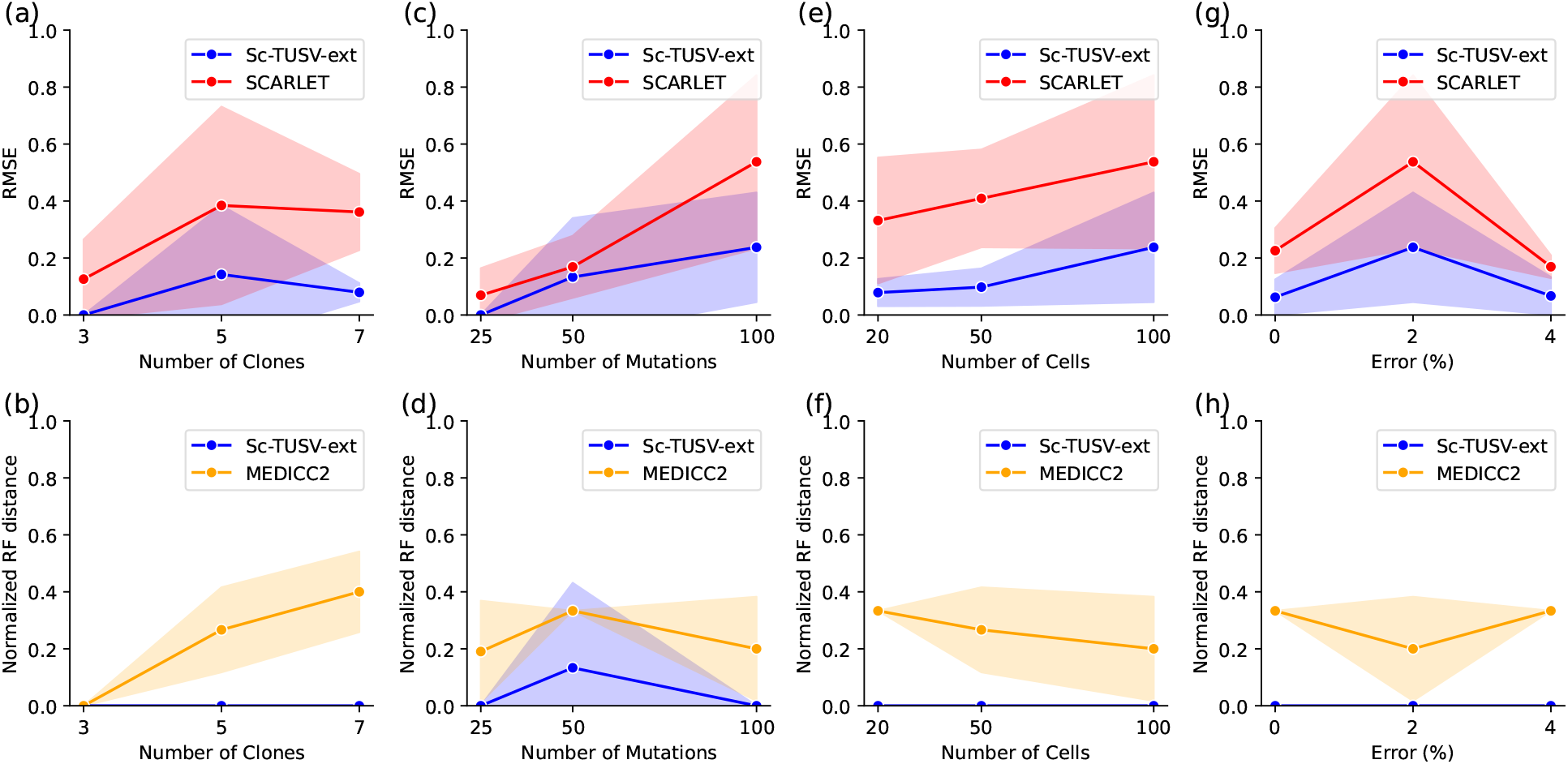
Results on experiment 1, 2, 3 and 4, with different number of clones, mutations and single-cells. (a), (c), (e) and (g) shows root mean squared error of the estimated SNVs with Sc-TUSV-ext and SCARLET with different model conditions; (b), (d), (f) and (h) shows the normalized RF distance of the inferred phylogenies with Sc-TUSV-ext and MEDICC2 for different model conditions. The solid lines represent the means of the simulations and shaded area represents one standard deviation on either side of the mean.

**Fig. 3.**
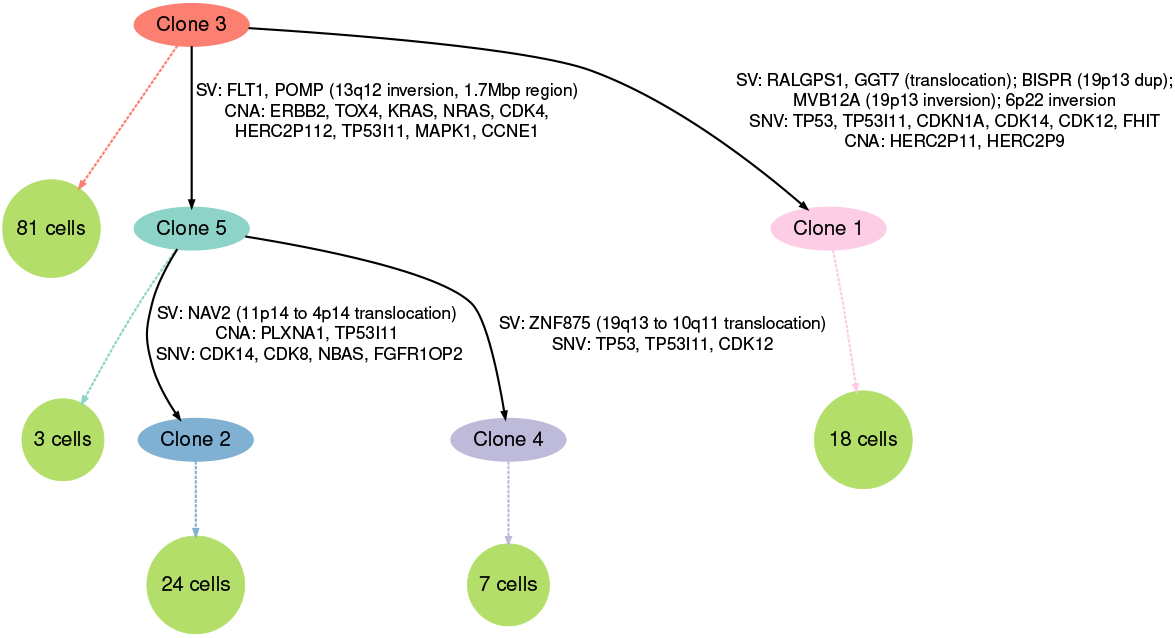
Sc-TUSV-ext inferred tree on DG1134 patient data. The edge labels represent subsets of mapped SNVs, CNAs and SVs. The number of cells mapped to the clones are shown with the dotted arrows. We subsampled to 120 SNVs and SV breakpoints and ran sc-TUSV-ext with 5 random restarts and 5 iterations for t = 5000.

To compare the output phylogeny of SC-TUSV-ext with the true phylogeny, we used the common ancestor set (CASet) distance as described in DiNardo et al. (8). For all cases, Sc-TUSV-extyielded a CASet distance = 0.0, which indicates Sc-TUSV-ext’s clonal phylogenies exactly resembles the true phylogeny’s. In addition, we compared the trees inferred by Sc-TUSV-ext and MEDICC2 with the true clonal trees (2(h)). To compare the output single-cell trees of MEDICC2 to the true and Sc-TUSV-ext’s clonal trees, we first substitute single-cells in the leaves of MEDICC2 trees with their clone assignments. We then transform Sc-TUSV-ext’s and the true clonal phylogenies to have the clones only in the leaves. We use normalized RF distance (28) of both MEDICC2 and SC-TUSV-ext phylogenies for different numbers of clones. We do not show a result for SCARLET, as we used SCARLET with a setting where the true tree was known. While we found accurate clusters with the MEDICC2 distances, the tree structure of MEDICC2 differed from the true one. In contrast, Sc-TUSV-ext provided us with accurate trees for different levels of error, notably due to the use of all types of variants, particularly SNVs.

### 3.3 Experiment 2: Validation for different number of clones

We tested our method with 3, 5, and 7 clones. For each case, we ran 5 simulations with 100 single cells, 100 SNVs and 100 SV breakpoints, 2% error in the SNVs, and 4% error in the CNAs. We compared the root mean squared error of our estimated SNVs with SCARLET’s (30) (Figure 2(a)). For each case, we ran Sc-TUSV-ext with a subsampling that considers 80 SV breakpoints and 40 SNVs. The results demonstrate that our method can estimate the SNVs more accurately than SCARLET for higher number of clonal populations. For each case, we first used MEDICC2 distances to cluster the single cells and found the adjusted Rand index (ARI) values of the single-cell clusters to be 1 and CASet distances of our estimated phylogenies to be 0 with respect to the true clusters and phylogenies for all 3, 5, and 7 clones. Similar to section 3.2, we noticed higher RF distances with MEDICC2-inferred phylogenies than those of Sc-TUSV-ext.

### 3.4 Experiment 3: Validation on different number of mutations

Next, we varied the number of mutations on a 5-clone tree with 100 single-cells and 2% error in the SNVs and 4% error in the CNAs. We varied the total number of SNVs, *m*_snv_ ∈ {25, 50, 100}, SV breakpoints, *m*_sv_ ∈ {50, 100, 200} and corresponding CNAs across the cells. Figure 2(c) demonstrates the RMSE of the estimated SNVs of SCARLET and Sc-TUSV-ext, and Figure 2(d) demonstrates the RF distances of the MEDICC2 phylogenies and Sc-TUSV-ext phylogenies with respect to the true phylogenies. We noticed the similar trend as before for Sc-TUSV-ext, SCARLET and MEDICC2. For 50 SNV mutations, one of our five replicates had a relatively low ARI (0.02) for the clusters which resulted in a tree with normalized RF distance = 0.67, explaining the high noise in the figure.

### 3.5 Experiment 4: Validation on different number of single-cells

Finally, we validated the robustness of our method by varying the number of single-cells, *N* ∈ {20, 50, 100}, with a 5 clone tree, 100 SNVs, 100 SV breakpoints, 2% error in the SNVs and 4% error in the CNAs. Figure 2(e) and 2(f) demonstrates the RMSE and RF distance between Sc-TUSV-ext, SCARLET and MEDICC2, which we found to be consistent with the conclusions of experiments 1, 2, and 3.

### 3.6 Application of Sc-TUSV-ext on real cancer data

We applied Sc-TUSV-ext to single-cell DNA sequencing (scDNAseq) data from a high-grade serous ovarian cancer (HGSC) cohort from Funnell et. al. (12). From the 16 HGSC cases, we chose one representative case, DG1134 (HGSC), as a demonstration of the performance of our method on real data. For this case, there were 133 single cells with structural variants that included fold back inversions (FBI) and as a result, high copy numbers. Funnell et al. derived single-cell copy numbers, SVs and SNVs using DLP+ (18), and used BWA-MEM (20) to align DLP+ reads to hg19 reference genome. The genomes were segregated into 500 kb bins. We further downsampled the data to run with MEDICC2 given our computational resources. We experimented with four separate downsampling models — (1) 5 Mbp bins, (2) 10 Mbp bins, (3) 20 Mbp bins, and (4) 500 Kbp bins with variance *>*= 3. We compared the ARI of our clones with the clones reported in the original publication (12) and found the highest ARI = 0.27 for 5 clones with (4). Apart from that, as FBI tumors from the original paper are reported to have a 1.9-fold higher copy number variance than other type of tumors, we ran MEDICC2 with copy number segments which have variance *>*= 3 across the single cells. Next, we chose one cell at random to represent each clone. Since there were many more SNVs than CNAs and SVs, we kept all the CNAs and SVs with numbers of SNVs set to at most 120 and mapped all the remaining SNVs to the tree afterwards. We ran Sc-TUSV-ext with 5 clones, 5 iterations, 5 random restarts and 5000 seconds per iteration on the initial data with 500 Kbp bins for CNAs.

Figure 3.6 shows Sc-TUSV-ext’s clonal phylogeny for DG1134, exhibiting a number of mutations in common driver genes (5). We found high amplification of CCNE1 in clones 2, 4 and 5, with copy numbers 2 in one allele and 9 in the other allele, consistent with the primary paper’s analysis of high amplification in that gene, although it is a subclonal mutation not repeated on the branch to clone 1. Similarly, the phylogeny shows one cell lineage (beginning with clone 5) exhibiting KRAS CNA mutation and FLT1 SV mutation, while a distinct lineage (beginning with clone 1) shows TP53 SNV mutation. These observations highlight the ability of our method to draw out distinct lineages acting through different driver pathways impacted by distinct mutation types. In addition, we found both SNV- and CNA-driven mutations in the TP53I11 gene in all the branches. These observations show the ability of our method to identify recurrent driver mutation, potentially indicative of strong selection, acting through multiple variant types. All branches of the phylogeny showed structural variations, indicating ongoing genomic instability shaping the tumor’s clonal evolution.

## 4 Discussion

In this work, we present Sc-TUSV-ext, a novel method for constructing clonal lineage trees (“tumor phylogenies”) from single-cell DNA sequencing data, while including SNVs, CNAs and SVs in the reconstruction. On simulated data, we showed the synergy of these distinct variant types in accurately resolving phylogenies, with Sc-TUSV-ext performing better both in estimating error-free variants and reconstructing accurate phylogenies than leading competitor methods that use only subsets of these variant types. We further demonstrated the value of integrating all of these variant types by applying Sc-TUSV-ext to an scDNA-seq data set from a high-grade serous ovarian cancer patient (12). The case showed how these different mechanisms of mutability can act in parallel in a single patient and how Sc-TUSV-ext’s comprehensive model can reveal important features of the biology that would be missed by other methods. There are, however, several directions by which Sc-TUSV-ext could be further improved. Sc-TUSV-ext currently relies on MEDICC2 for clustering the single-cells, limiting its ability to capture fully high (*>*= 8) copy number changes. It also currently lacks ability to handle some classes of complex complex structural variations, such as chromothripsis or chromoplexy (2). It would also benefit from improved scalability to allow it to make better use of large variant sets.

## 5 Acknowledgements

This work was supported in part by a gift from the Mario Lemieux Foundation. Research reported in this publication was supported by the National Human Genome Research Institute of the National Institutes of Health under award number R01HG010589. The content is solely the responsibility of the authors and does not necessarily represent the official views of the National Institutes of Health.

